# Brain orchestration of pregnancy and maternal behavior in mice

**DOI:** 10.1101/2020.05.23.112045

**Authors:** David André Barrière, Arsène Ella, Frédéric Szeremeta, Hans Adriaensen, William Même, Elodie Chaillou, Martine Migaud, Sandra Même, Frédéric Lévy, Matthieu Keller

## Abstract

Reproduction induces changes within the brain to prepare for gestation and motherhood. However, the dynamic of these central changes and their relationships with the development of maternal behavior remain poorly understood. Here, we describe a longitudinal morphometric neuroimaging study in female mice between pre-gestation and weaning, using new magnetic resonance imaging (MRI) resources comprising a high-resolution brain template, its associated tissue priors (60-μm isotropic resolution) and a corresponding mouse brain atlas (1320 regions of interest). Using these tools, we observed transient hypertrophies not only within key regions controlling gestation and maternal behavior (medial preoptic area, bed nucleus of the *stria terminalis*), but also in the amygdala, caudate nucleus and hippocampus. Additionally, unlike females exhibiting lower levels of maternal care, highly maternal females developed transient hypertrophies in somatosensory, entorhinal and retrosplenial cortices among other regions. Therefore, coordinated and transient brain modifications associated with maternal performance occurred during gestation and lactation.

## Introduction

Motherhood is among the most transformative experiences in the lives of female mammals. While virgin females tend to avoid neonates, the end of the gestation period and the birth process lead to a behavioral switch characterized by an attraction towards infant cues, the expression of nurturing behavior and ultimately the establishment of infant bonding^1,2^. Decades of scientific research dedicated to the maternal brain have revealed a core neural circuitry that includes the medial preoptic area (mPOA) and the adjoining ventral part of the bed nucleus of the *stria terminalis* (BNSTv), and that is highly critical for the onset of maternal behavior^1,3–5^. Functional modulations of the mPOA/BNSTv consistently disrupt maternal motivation and expression in numerous species^1,2,6^. This core maternal circuitry regulates maternal behavior through its direct projections to the ventral tegmental area, which promotes reward system activation^1,7^, as well as through its connections with cortical regions, including the prefrontal cortex^8–10^. This crucial central circuitry is finely regulated by multiple neural networks that integrate both internal and external stimulations. The proper expression of maternal care towards offspring is prepared through the neuroendocrine action of sex steroids and neuropeptides such as oxytocin among others during the gestation period^11^. These internal factors induce rewiring of the maternal brain, including through structural plasticity through increasing neuronal soma size or astrocytic complexity within the mPOA^12^, and changes in neurogenesis mainly in the main olfactory bulb (MOB)^13^ but also in the mPOA/BNST in rodents^14^. In human, regional morphological changes of gray matter (GM) within the parahippocampal gyrus, precuneus, cingulate, insula and frontal cortex have been observed in primiparous women using magnetic resonance imaging (MRI)^15^. Additionally, olfactory cues coming from the neonate are integrated by the MOB and the accessory olfactory bulb (AOB) through an amygdalo-hypothalamic pathway, which is responsible for attraction/repulsion behavioral outcomes^16^. Hence, the development of the maternal brain is dependent on both integration of external and internal cues acting through multiple brain pathways and regions to prepare the brain to gestation and motherhood. Nevertheless, the relationship between the brain rewiring over the gestation and lactation periods and the establishment of the maternal behavior is poorly documented.

To assess the dynamics of the maternal brain, a longitudinal MRI morphometric study over a complete reproductive experience was performed in mouse to investigate changes in the gray matter concentration (GMC) using voxel-based morphometry (VBM). VBM is a well-established and well-validated image analysis technique that provides an unbiased and comprehensive assessment of anatomical differences throughout the brain, and has been successfully used to study GM changes within the mouse brain^17–21^. However, available mouse brain MRI resources are often partial or provided in different spatial orientations or spatial resolutions (**Table 1**). As an example, the <u>Australian Mouse Brain Mapping Consortium</u> (AMBMC), offers a high resolutive template and detailed atlases of the mouse brain including the cerebellum^22^, hippocampus^23^, diencephalon^24^ and cortices^25^. Unfortunately, segmentation of the MOB and the AOB and hindbrain is lacking, and this resource does not provide associated tissues probabilistic maps necessary for VBM. In other hand, the <u>Allen Mouse Brain Common Coordinate Framework</u>, is the most advanced mouse brain atlas^26^. This new atlas delimitates discrete structures within the thalamus, hindbrain, olfactory system and diencephalon and provides a full segmentation of cortical layers however, MRI template and brain tissues priors are still lacking for VBM investigations.

**Table 1.** Comparison of mouse brain resources currently available in the literature (NA = not available).

| *Ex vivo /*<br>*In vivo* | Strain | Sex | Age<br>(days) | Number of<br>animals | Magnetic field<br>intensity | Anatomical<br>contrast | Spatial<br>resolution | MRI<br>template | Atlas | Regions<br>of interest | Tissue<br>Probability<br>maps | References |
| --- | --- | --- | --- | --- | --- | --- | --- | --- | --- | --- | --- | --- |
| *Ex vivo* | C57BL/6J | Male | 63 | 6 | 9.4 Tesla | T<sub>1</sub><br>PD<br>T<sub>2</sub>W | 90 x 90 x 90<br>μm<sup>3</sup> | Yes | Yes | 21 | No | Ali et al.<br>2005 |
| *In vivo* | C57BL/6J | Male | 100 | 6 | 11.7 Tesla | T<sub>1</sub> | 60 x 60 x 60<br>μm<sup>3</sup> | Yes | Yes | NA | No | MacKenzie-Graham et<br>al. 2011 |
| *Ex vivo* | 129S1/SvImJ | Male | 56 | 9 | 7 Tesla | T<sub>2</sub> | 60 x 60 x 60<br>μm<sup>3</sup> | Yes | Yes | 9 | No | Kovachev et al. 2005 |
| *Ex vivo* | C57BL/6J | NA | PO | 8 | 11.7 Tesla | T<sub>1</sub> | 40 x 40 x 40<br>μm<sup>3</sup> | Yes | Yes | 12 | Yes | Lee et al.<br>2005 |
| *In vivo* | C57BL/6J | Male | 84-98 | 12 | 9.4 Tesla | T<sub>2</sub> | 100 x 100 x 100<br>μm<sup>3</sup> | Yes | Yes | 20 | No | Ma et al.<br>2008 |
| *In vivo* | C3H/HeSnJ | NA | 77 | 15 | 7 Tesla | T<sub>1</sub> | 156 x 156 x 156<br>μm<sup>3</sup> | Yes | Yes | 6 | No | Brack et al.<br>2006 |
| *Ex vivo* | 129S1/SvImJ<br>C57BL6<br>CTD1 | Male | 126 | 27 | 7 Tesla | T<sub>2</sub> | 60 x 60 x 60<br>μm<sup>3</sup> | Yes | Yes | 42 | No | Chen et al.<br>2008 |
| *Ex vivo* | C57BL/6J | NA | 63 | 6 | 9.4 Tesla | T<sub>1</sub><br>T<sub>2</sub> | 21.5 x 21.5 x 21.5<br>μm<sup>3</sup><br>43 x 43 x 43<br>μm<sup>3</sup> | Yes | Yes | 33 | No | Badea et al.<br>2008 |
| *Ex vivo* | C57BL/6J | 20 males<br>20 females | 84 | 40 | 7 Tesla | T<sub>1</sub> | 32 x 32 x 32<br>μm<sup>3</sup> | Yes | Yes | 62 | No | Dorr et al.<br>2008 |
| *Ex vivo* | C57BL/6J<br>and<br>BXD | Male | 63 | 12 | 9.4 Tesla | T<sub>1</sub><br>T<sub>2</sub> | 21.5 x 21.5 x 21.5<br>μm<sup>3</sup><br>43 x 43 x 43<br>μm<sup>3</sup> | Yes | Yes | 20 | No | Sharief et al.<br>2008 |
| *Ex vivo* | C57BL/6J | Male | 66-78 | 14 | 9.4 Tesla | T<sub>1</sub><br>T<sub>2</sub><br>T<sub>2</sub>* | 21.5 x 21.5 x 21.5<br>μm<sup>3</sup> | Yes | Yes | 37 | No | Johnson et al.<br>2010 |
| *Ex vivo* | C57BL/6J | Male | 84 | 18 | 16.4 Tesla | T<sub>1</sub>/T<sub>2</sub>* | 30 x 30 x 30<br>μm<sup>3</sup> | Yes | Yes<br>(Partial:<br>hippocampus) | 40 | No | Richards et al.<br>2011 |
| *Ex vivo* | C57BL/6J | Male | 84 | 18 | 16.4 Tesla | T<sub>1</sub>/T<sub>2</sub>* | 30 x 30 x 30<br>μm<sup>3</sup> | Yes | Yes<br>(Partial:<br>cerebellum) | 38 | No | Ullman et al.<br>2013 |
| *Ex vivo* | C57BL/6J | Male | 84 | 18 | 16.4 Tesla | T<sub>1</sub>/T<sub>2</sub>* | 30 x 30 x 30<br>μm<sup>3</sup> | Yes | Yes<br>(Partial: basal<br>ganglia) | 35 | No | Ullman et al.<br>2013 |
| *Ex vivo* | C57BL/6J | Male | 84 | 18 | 16.4 Tesla | T<sub>1</sub>/T<sub>2</sub>* | 30 x 30 x 30<br>μm<sup>3</sup> | Yes | Yes<br>(Partial:<br>neocortex) | 74 | No | Ullman et al.<br>2013 |
| *In vivo* | C57BL/6J | Male | 126 | 82 | 4.7 Tesla | T<sub>2</sub> | 70 x 70 x 70<br>μm<sup>3</sup> | No | NA | NA | Yes | Sawak et al.<br>2013 |
| *Ex vivo* | C57BL/6J | Male | 84 | 18 | 16.4 Tesla | T<sub>1</sub>/T<sub>2</sub>* | 30 x 30 x 30<br>μm<sup>3</sup> | Yes | Yes<br>(Partial:<br>diencephalon) | 89 | No | Watson et al.<br>2017 |
| *Ex vivo* | C57BL/6J | 1,051 males<br>621 female | 77 | 1675 | NA | NA | 10 x 10 x 10<br>μm<sup>3</sup> | No | Yes | 1,327 | No | Wang et al.<br>2020 |

Thus, we combined these resources and associated probabilistic maps to emulate a complete resource dedicated to the mouse brain. Using these resources and a longitudinal VBM approach, we were able to assess the dynamic morphological changes of the brain during the whole reproductive period and demonstrate how these changes predict the quality of maternal behavior.

## Results

### Mouse MRI atlas

VBM strategies require a template image and its associated priors of gray matter (GM), white matter (WM) and cerebrospinal fluid (CSF) for brain segmentation and normalization. In addition, a complete atlas of the mouse brain is mandatory for the identification of regions of interest (ROIs) highlighted by the VBM analysis. Given the limitations of available tools to thoroughly study GMC changes during the gestation and lactation periods in mice, we developed first a new set of resources using the AMBMC, an ultra-high-resolution template built from *ex vivo* brain images finely normalized within the same space^25^ and the <u>Allen Mouse Brain Common Coordinate Framework</u>^26^ (**Figure 1**). Our resources comprise the following four components: 1) a complete mouse brain template with a spatial resolution suitable for mouse brain analysis (60-μm isotropic resolution); 2) the corresponding GM, WM and CSF probabilistic maps for brain normalization together with a VBM analysis built from 138 T_2_-weighted images; 3) a complete mouse brain atlas derived from Paxinos and Franklin’s mouse brain atlas^27^ and composed of a mosaic of 1320 ROIs (**Figure 2A**); 4) a brain mesh permitting brain plot generation and data visualization (**Figure 2B** and **Supplemental Video 1**). We visually inspected and carefully checked the results of the normalization process against the original coregistered atlas. Then, the labeled structures were reclassified and aggregated according to the brain regions to which they belonged (auditory, insular, temporal cortices, etc.), with respect to their anatomical topography (cortex, basal ganglia, etc.), tissue type (GM, WM and CSF) and hemisphere (left or right). Cortical structures were subdivided into functional (*e.g.*, primary and secondary motor cortices) or structural (agranular, dysgranular, agranular/dysgranular, granular and posterior agranular insular cortices) areas (**Figure 2C**), as well as into different cortical layers (**Figure 2D**). Subcortical structures, as for example, the hypothalamus (**Figure 2E**) and the hippocampus (**Figure 2F**), were fully segmented according to Paxinos and Franklin’s atlas^27^.

**Figure 1.**
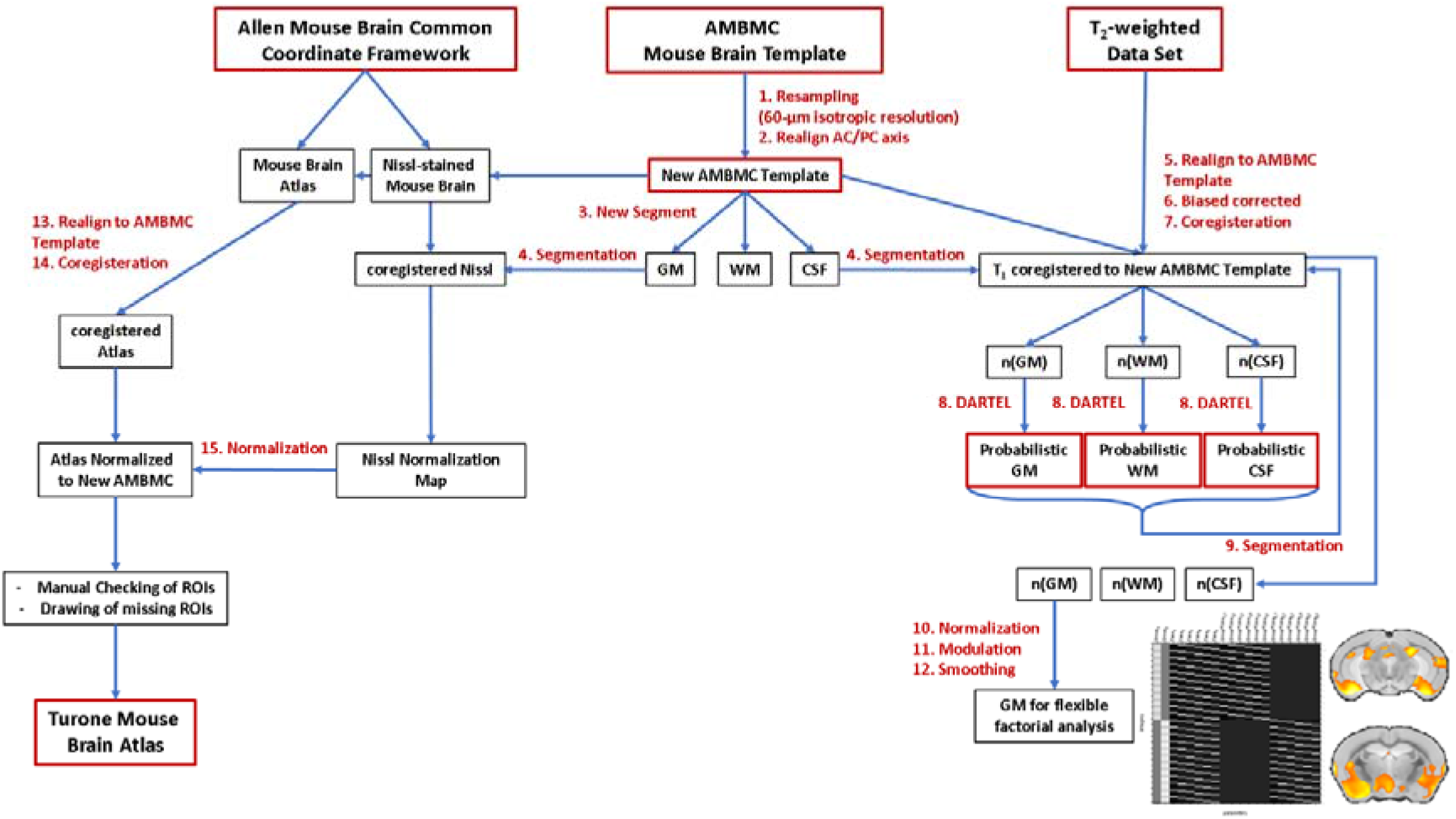
Processing of mouse brain templates and building an atlas from the AMBMC template and Allen Brain Atlas for data analysis and visualization. To create our resources, we used both AMBMC mouse brain template and the mouse Allen Brain Atlas and its associated Nissl images. (**1**) We down-sampled to a suitable resolution for MRI analysis (60-μm isotropic resolution) and (**2**) realign in the AC/PC axis. The resulting template was then segmented into GM, WM and CSF probability maps (**3**). These probability maps were used to segment all the images which have been previously normalized to the template (**5**,**6**,**7**). We obtained a large set of 138 images for each tissue class which have been used to build a population-specific GM, WM and CSF priors. using an exponentiated lie algebra (DARTEL) approach (**8**). This new set of population-specific tissue priors was used to segment again normalized T_2_ images (**9**) for the final VBM preprocessing (**10**, **11**, **12**). To normalize the Allen Brain Atlas, we manually realign (**13**) and normalized the associated Nissl-stain mouse brain using the GM priors generated previously (**14**, **15**). Both linear and nonlinear transformations have been applied to the Allen mouse brain atlas. Then, a visual inspection of each normalized label was carried out and, when necessary, redrawn according to Paxinos and Franklin’s atlas. Finally, the olfactory bulb and hind brain regions were completed, the *corpus callosum* and ventricles were drawn from the WM and CSF priors, and the cerebellum labels were replaced by the AMBMC cerebellum labels. AMBMC = Australian Mouse Brain Mapping Consortium mouse brain template, AC/PC = anterior commissure/posterior commissure, CSF = cerebrospinal fluid, GM = gray matter, WM = white matter.

**Figure 2.**
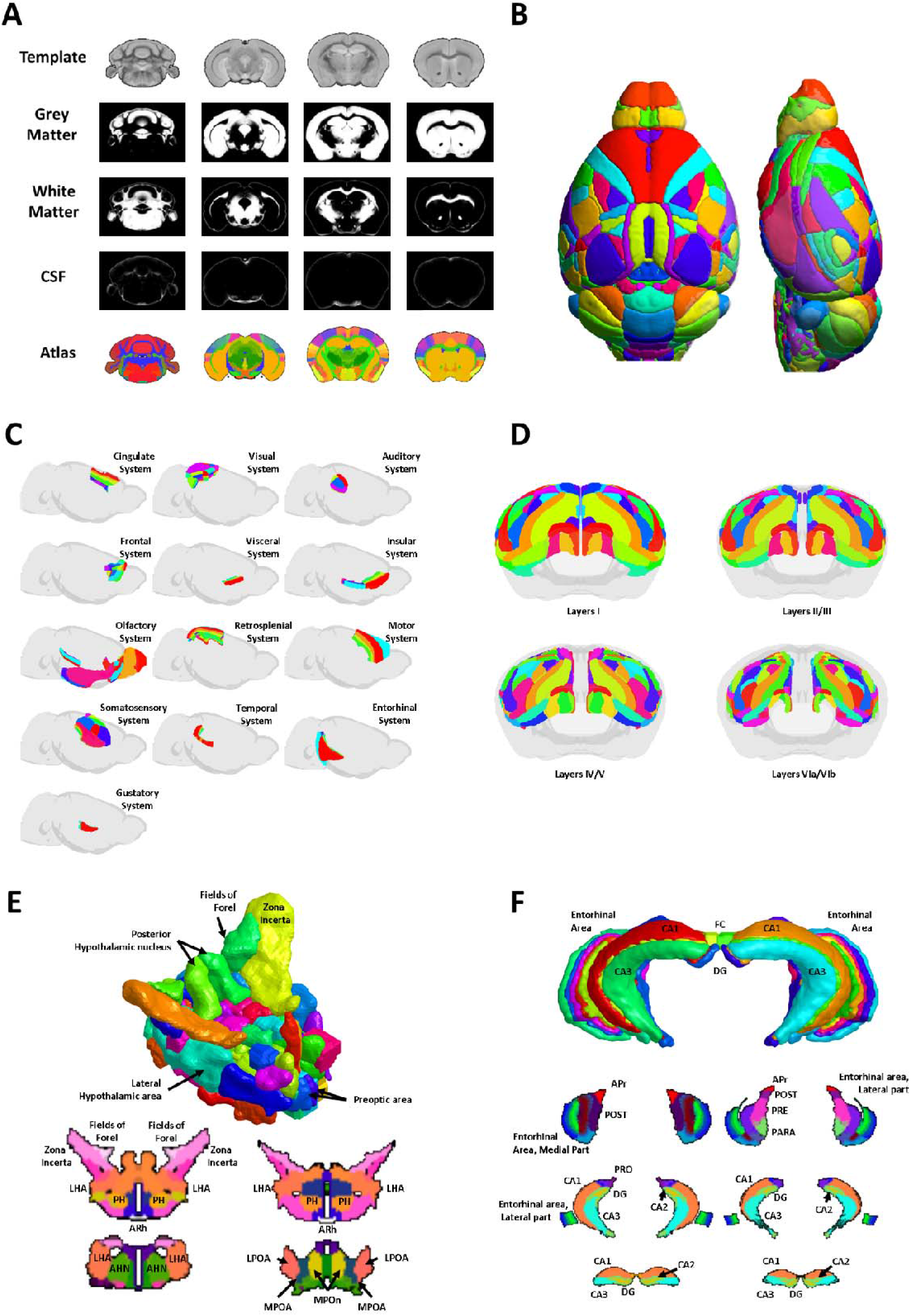
Details of the mouse brain template and atlas. (**A**) Coronal slices of the anatomical template of the mouse brain and the corresponding gray matter, white matter and cerebrospinal fluid probabilistic maps and the associated anatomical atlas (60-μm isotropic resolution). (**B**) Dorsal (left panel) and lateral (right panel) 3D representations of the anatomical mouse brain atlas. (**C**) Lateral views of the cortical areas after normalization of the Allen Mouse Brain Atlas to the AMBMC anatomical template. The cortex was segmented into cortical areas such as the cingulate, visual, auditory, frontal, visceral, insular, olfactory, retrosplenial, motor, somatosensory, temporal, entorhinal and gustatory systems. Each area was subdivided into secondary areas (*e.g.*, primary and secondary motor cortices) or structural areas (*i.e.*, agranular, dysgranular, agranular/dysgranular, granular and posterior agranular insular cortices). (**D**) The four images depict the different cortical layers (I, II/III, IV/V and VIa/VIb). (**E** and **F**) 3D rendering and axial sections of subcortical structures (hypothalamus and hippocampus). Legend for labeled regions: <u>Hypothalamus</u>: *ARh = arcuate hypothalamic nucleus; LHA = lateral hypothalamic area; LPOA = lateral preoptic area; MPOA = medial preoptic area; MPOn = medial preoptic nucleus; PH = posterior hypothalamic nucleus. AHN = anterior hypothalamic nucleus*. <u>Hippocampus</u>: *Apr = area prostriata; CA1, CA2, CA3 = cornu ammonis areas 1, 2 and 3; DG = dentate gyrus; FC = fasciola cinerea; PARA = parasubiculum; POST = postsubiculum; PRO = prosubiculum; PRE = presubiculum.* For further details, see https://www.nitrc.org/projects/tmbta_2019.

### Morphometric changes occurred during the gestation and lactation periods

Next, we used our new resources to study the variations of GMC in mouse brain from the beginning of gestation until weaning. Using MRI T_2_-weighted anatomical acquisitions, we estimated the GMC maps which offer for each animal a global estimation of the GM. Longitudinal comparison of GMC between virgin mice (control group, n= 11) and mice who became pregnant and raised their young until weaning (parous group, n=12) permits to highlight local modifications of GM during the whole reproductive cycle. A comparison of baseline and early gestation GMC maps between the control and parous groups did not reveal significant differences. However, at the end of the gestation period, significant increases in GMC were observed within several brain regions in the parous group compared to the control group (**Table S1** and **Figure 3A**). Time course analysis revealed differences in GMC profiles between control and parous groups at the end of the gestation period, early in lactation and at the end of the lactation period. Specifically, GMCs within the mPOA and the BNST were consistently significantly higher in the parous group at that times. In addition, within the agranular insular cortex in the late gestation period and the early lactation period, GMC was significantly higher in parous group compare to control group (**Figure 4A**). During the early lactation period, we also found specific and significant increases in GMCs of the parous group within numerous brain regions (**Table S2** and **Figure 3B**). Among these structures, the hippocampus (CA1 layer), amygdalar area and piriform area showed a transient increase in GMC at the early lactation time point that returned to baseline values at the end of the lactation period in parous group compared to control group (**Figure 4B**). In contrast, the caudate putamen, arcuate nucleus and paraventricular nucleus of the hypothalamus (PVN) showed significantly higher GMCs in the parous mice than in the control mice during the lactation period (**Table S3** and **Figures 3C and 4C**). Together, our data demonstrate that the late gestation period is associated with a pronounced increase in GMC in the mPOA/BNST, the core neural system of maternal motivation, lasting up to the late lactation period. Furthermore, the early lactation period is associated with increased GMC in other key maternal motivation areas in midbrain regions, including the hypothalamus, caudate putamen and amygdala. These GMC differences between both groups were no longer observed after weaning.

**Figure 3.**
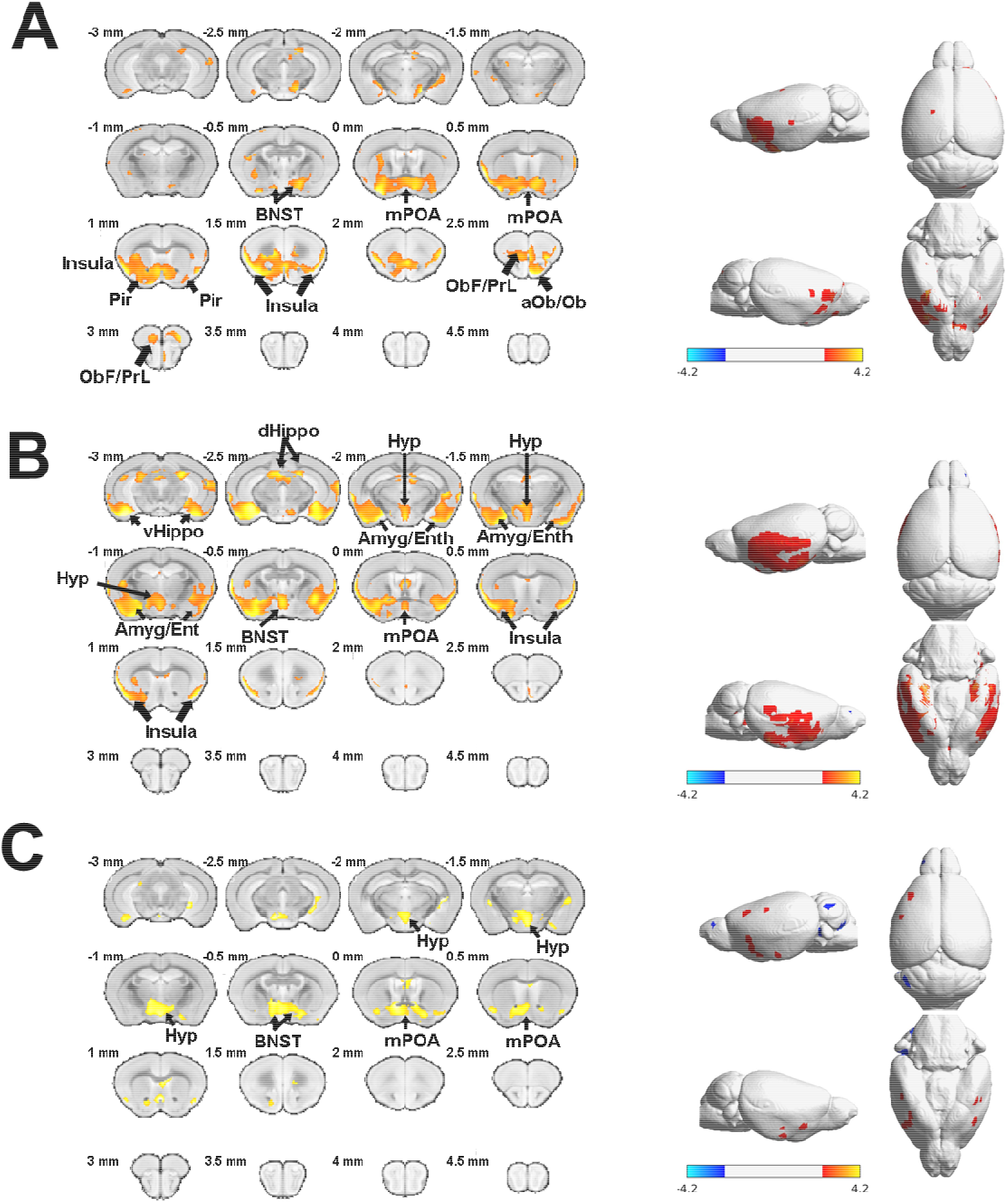
Longitudinal effects of the reproductive cycle on brain morphometry. Coronal slices (at left) and brain plots (at right) showing gray matter concentration (GMC) differences between control and parous animals at the end of gestation (**A**) and during early lactation (**B**) and late lactation (**C**). SPM flexible factorial analysis revealed an interaction between control mice and parous mice in the late gestation period (**A**), early lactation period (**B**) and late lactation period (**C**); voxel-level threshold *p* < 0.01, t_(126)_ = 2.356, cluster threshold = 25 voxels. *BNST* = *bed nucleus of the stria terminalis*; *Hyp* = *hypothalamus*; *mPOA* = *medial preoptic area*; *dHippo* = *dorsal hippocampus*; *ObF/PrL* = *orbitofrontal/prelimbic area*; *aOb/Ob* = *accessory olfactory nucleus/olfactory bulb*; *Pir* = *piriform cortex*; *Amyg/Ent* = *amygdala/entorhinal cortex*.

**Figure 4.**
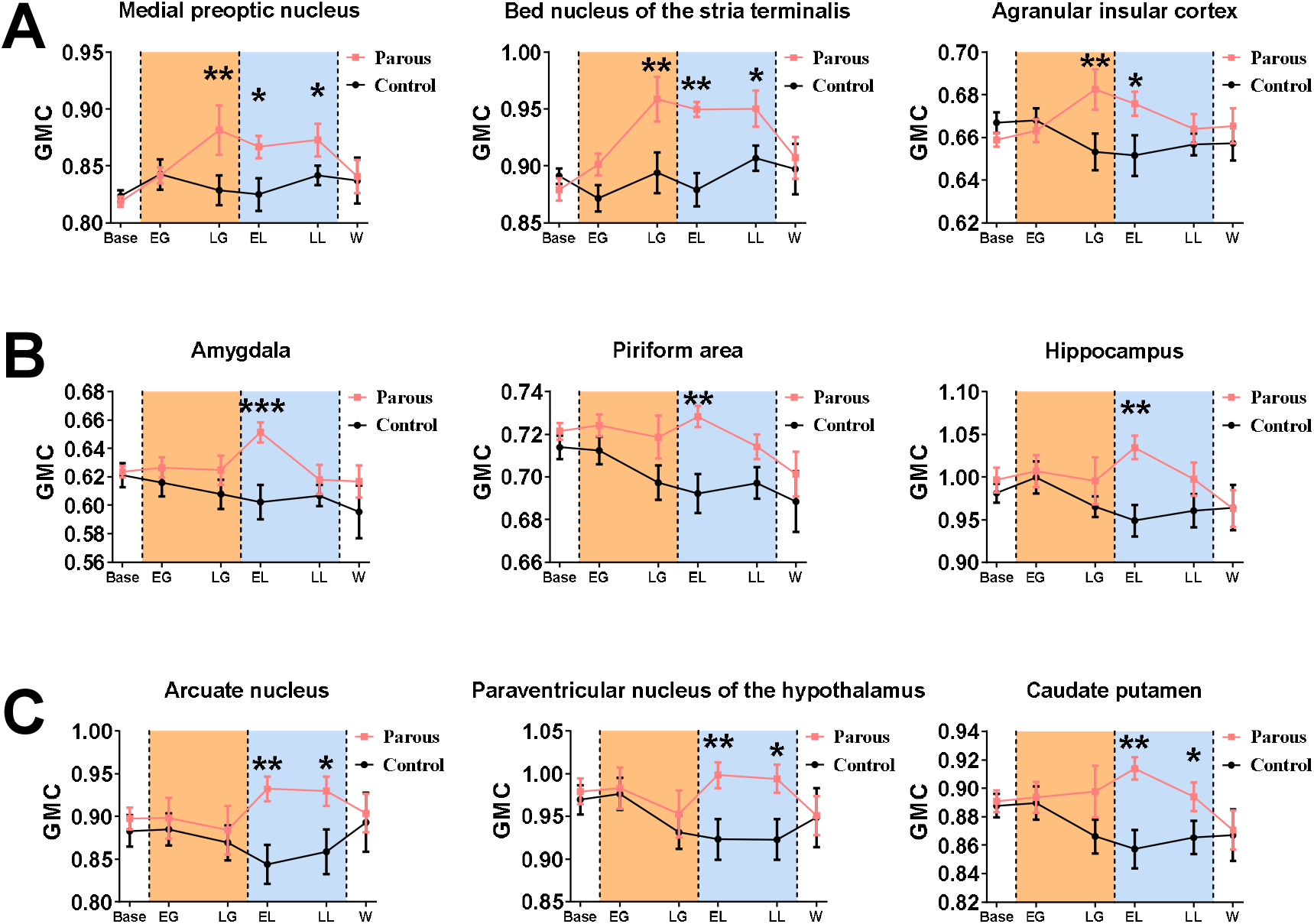
Longitudinal time course analysis of gray matter concentration (GMC) over the reproductive cycle. Time course comparisons in GMC between the control (black dots and lines) and parous (red dots and lines) groups showing 3 different time profiles. (**A**) GMC values within the medial preoptic area, the bed nucleus of *stria terminalis* (BNST) and the agranular insular cortex reveal a significant increase in GMC during the late gestation (LG) period maintained until weaning (W). (**B**) Specific and transient increases in GMCs are observed in the amygdala, the piriform area and the hippocampus during early lactation (EL). (**C**) The arcuate nucleus, PVN and caudate putamen display an increase in GMC through both EL and late lactation (LL) periods. Orange and blue areas represent the gestation and lactation periods, respectively. Data are expressed as the mean ± standard error of the mean (SEM); two-way ANOVA followed by Holm-Sidak multiple comparisons test; \**p* < 0.05, ** *p* < 0.01 and *** *p* < 0.001, compared with control mice.

### Morphometric changes during gestation predict the quality of maternal behavior

In the last part of this work, we evaluated whether these morphometric changes might reflect differences in maternal performance. Based on 15 min of behavioral observation during the pup retrieval test performed one week after birth, we evaluated the maternal performance of each mother by measuring the first, second and third pup retrieval times, pup-licking duration, crouching time, rearing time, digging time and self-grooming time (**Figure 5A**). Interestingly, we observed a large distribution of values for both crouching (284.2 s, SD ± 268.7 s) and digging (98.22 s, SD ± 159.3 s) durations within the parous group. Whereas crouching duration relates to maternal behavior, digging duration is widely recognized as a discriminative marker of stress-related behavior in rodents^28^. Hence, we used crouching and digging durations to cluster animals using the k-means clustering procedure, thereby clustering parous animals into a high maternal behavior group (those with a high crouching time and low digging time) and a low maternal behavior group (those with low crouching time and high digging time).

**Figure 5.**
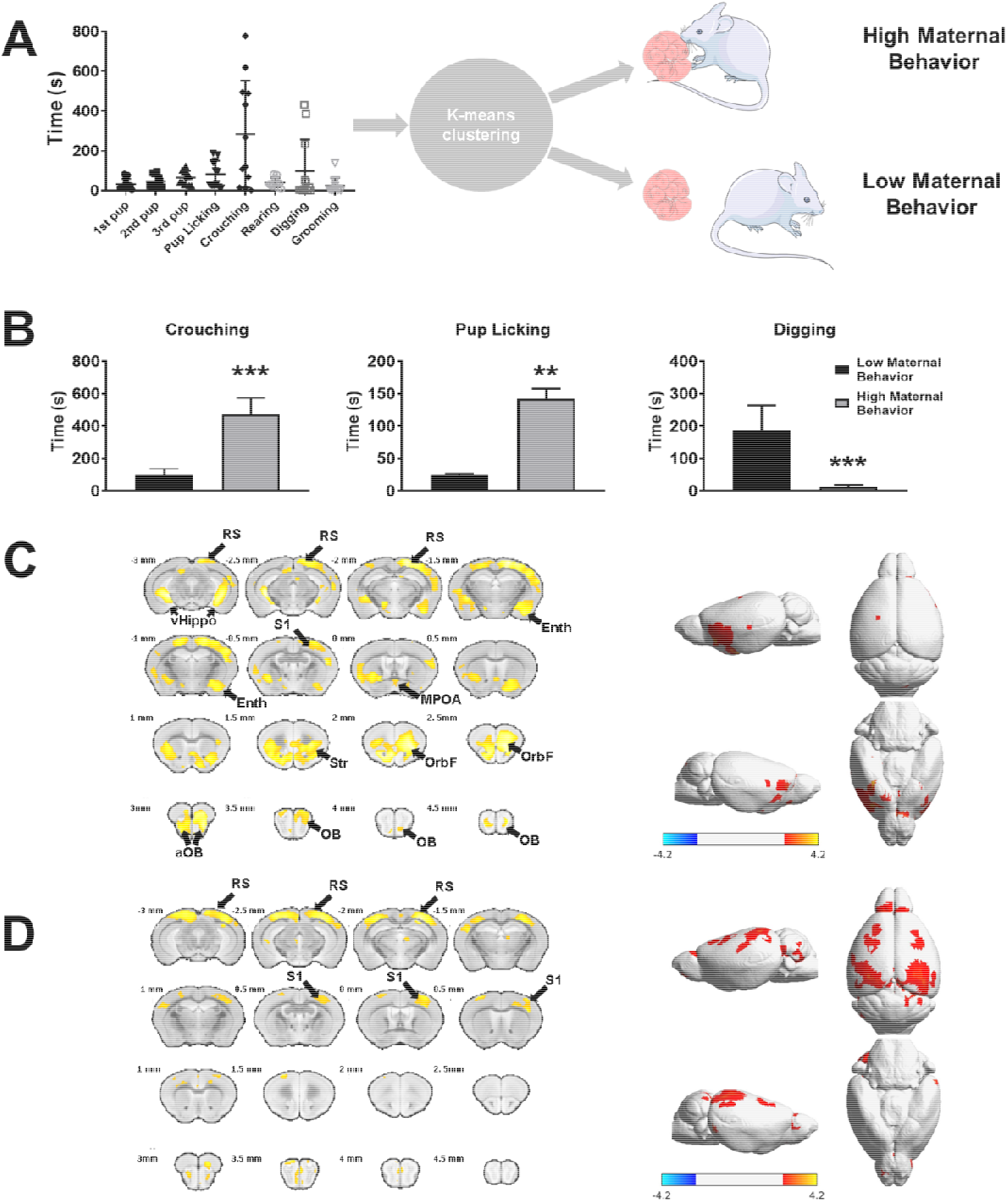
Distribution of animals according to the quality of their maternal behavior assessed with the pup retrieval test and brain morphometric. K-means clustering of parous animals to classify mice into low and high maternal behavior groups based on behavior during the pup retrieval test (**A**). Comparisons between the low and high maternal behavior groups revealed significant differences in crouching, pup-licking and digging times (**B**). Brain slices (left panel) and brain plots (right panel) comparing gray matter concentration (GMC) modifications and surface maps of GMC differences between females exhibiting low and those exhibiting high maternal behavior at the end of the gestation period (**C**) and early lactation period (**D**). Low and high maternal behavioral data were compared using a Student’s t-test with post hoc corrections for multiple comparisons using an FDR approach (Q = 1%) and are expressed as the mean ± SEM; ** *p* < 0.01 and *** *p* < 0.001. SPM flexible factorial analysis revealed an interaction between low and high maternal behavior parous mice in the late gestation period (**A**) and early lactation period (**B**); voxel-level threshold *p* < 0.01, t_(60)_ = 2.39, cluster threshold = 25 voxels. *RS* = *retrosplenial cortex*; *vHippo* = *ventral hippocampus*; *S1* = *primary somatosensory cortex*; *aOB* = *accessory olfactory bulb*; *OB* = *olfactory bulb*; *Pir* = *piriform cortex*; *Enth* = *entorhinal cortex*; *Str*= *striatum*; *OrbF* = *orbitofrontal cortex*.

A comparison of maternal performance parameters between the two clustered groups revealed significantly lower crouching and pup-licking times and a significantly higher digging time (**Figure 5B**) in the low maternal behavior group (n=6) than in the high maternal behavior group (n=6). Moreover, a comparison of GMC maps revealed both cortical and subcortical differences between the two clustered groups, mainly in the late gestation period (**Table S4** and **Figure 5C**) but also during the early lactation period (**Table S5** and **Figure 5D**). Indeed, transient increases in GMCs within the entorhinal area, lateral part of the orbital area, AOB, and medial preoptic area hat were observed at the end of pregnancy in the high maternal behavior group were absent in the low maternal behavior group (**Figure 6A**). In addition, the high maternal behavior group showed consistent higher GMCs in the hippocampus, retrosplenial area and barrel field of the primary somatosensory cortex (**Figure 6B**) from the end of the gestation until the end of the lactation period.

**Figure 6.**
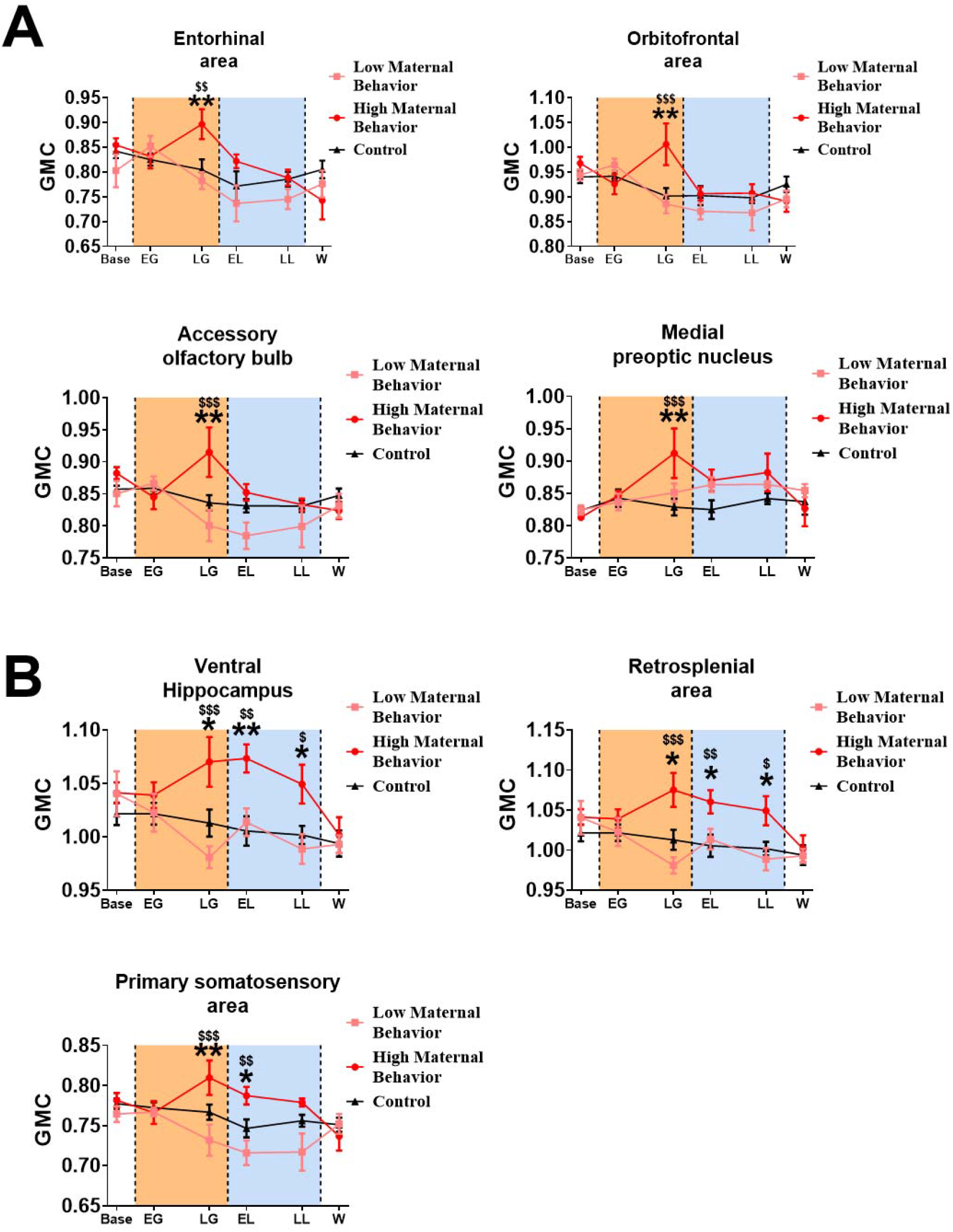
Longitudinal analysis in gray matter concentration (GMC) during the reproductive cycle in control females and females exhibiting low or high maternal behavior. Time-course comparison of GMC between the control (black dots and lines), low maternal behavior (pink dots and lines) and high maternal behavior (red dots and lines) groups revealed two types of time profile. GMC analysis in the entorhinal area, orbitofrontal area, the accessory olfactory bulb and the medial preoptic nucleus revealed an acute and specific increase in GMC values in the high maternal behavior group at the late gestation period (**A**). In contrast, GMC analysis in the ventral hippocampus, the retrosplenial area and the primary somatosensory area revealed an increase in GMC in the high maternal behavior group at the late gestation period, and this increase was maintained until weaning (**B**). Orange and blue areas represent the gestation and lactation periods, respectively. Data were compared using a two-way ANOVA followed by Holm-Sidak multiple comparisons test and expressed as the mean ± SEM; * *p* < 0.05, ** *p* < 0.01 when high maternal mice were compared with control mice and ^$$^ p <0.01, ^$$$^ p < 0.001 when high maternal mice were compared with low maternal mice.

Interestingly, using a receiver operating characteristic (ROC) analysis, we found that GMCs values within the entorhinal area and AOB at late gestation are reliable predictors for mouse maternal performance after birth. These GMC values significantly distinguished low maternal performance from high maternal performance postpartum (entorhinal area: sensitivity = 100, confidence interval (CI) = 61% to 100%; specificity = 83, CI = 44% to 99%; likelihood ratio = 6; **Figure 7A**; AOB: sensitivity =100, CI = 61% to 100%; specificity = 83, CI = 44% to 99%; likelihood ratio = 6, **Figure 7B**). The GMC values of both the entorhinal area and AOB observed at late gestation were also significantly correlated with maternal behavior (crouching and digging times). These results reveal that the GMC differences in olfactory (AOB and entorhinal cortex) and mnesic (entorhinal area)-related brain regions occurring during the late gestation period significantly predicted the quality of maternal behavior.

**Figure 7.**
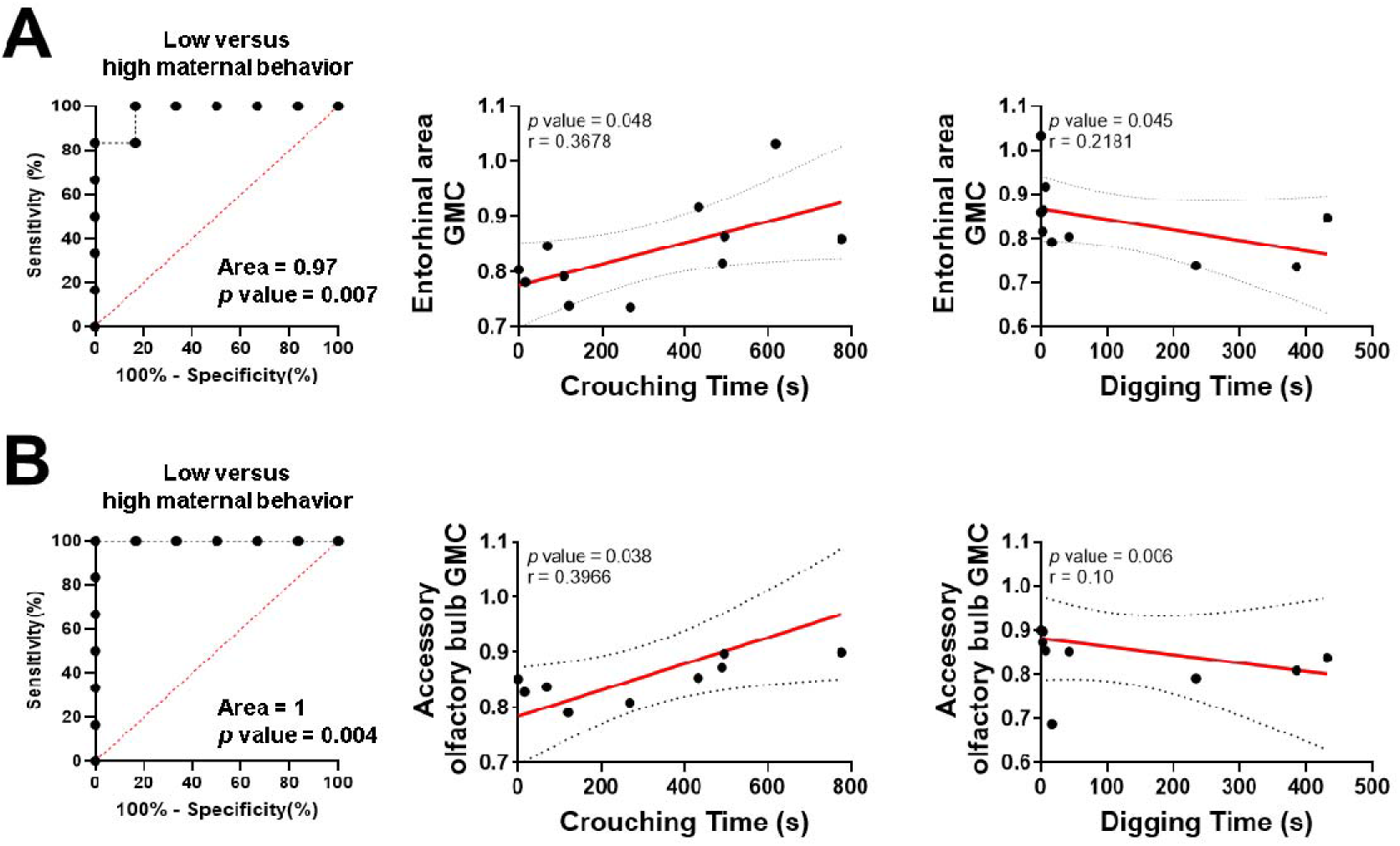
Estimation of the sensitivity and specificity of late-gestation GMC measures in the entorhinal area (**A**) and accessory olfactory bulb (**B**) to predict postpartum maternal performance. Receiver operating characteristic (ROC) curves were estimated using a Wilson/Brown test with a 95% confidence interval. Correlations were estimated using a Pearson correlation test. ROC and correlation analyses were considered significant at *p* < 0.05.

## Discussion

Using a new comprehensive neuroimaging resource dedicated to mouse brain, this longitudinal study reveals that pregnancy and lactation coincide with pronounced and transient cerebral changes. Transient increases in GMCs were observed in key regions controlling maternal behavior (mPOA, BNST, and PVN), as well as regions involved in emotions (amygdala), in motivation and reward (caudate nucleus, orbitofrontal cortex) and in mnesic functions (hippocampus). Interestingly, increase in GMC was also revealed in the insular cortex thought to link social and emotional skills. Moreover, we showed that females expressing high levels of maternal behavior had developed specific increases in GMCs in structures involved in olfactory (MOB and AOB) and somatosensory (somatosensory cortex) information processing, in memory (hippocampus, entorhinal cortex, retrosplenial cortex) and in reward and reinforcement (striatum). Interestingly, these hypertrophies were already significant at the end of the gestation period thus being predictive of the quality of maternal care (**Supplemental Video 2**).

### Implementation of new resources to support the analysis of mouse brain MRI data

The use of preclinical MRI is a target of growing interest for the study of brain structure and function in both healthy and pathological conditions. The use of advanced MRI techniques, coupled with the development of advanced animal models, is a powerful way to push new breakthroughs in the understanding of brain functioning and pathology. Herein, in the first step in our study, from the recent major advances in the development of brain mouse atlas we generated a new set of neuroinformatic tools offering for the first time a complete resource dedicated to MRI studies of the mouse brain, namely, an accurate brain atlas (1320 ROIs), a high-resolution brain template and the associated GM, WM and CSF priors (60-μm isotropic resolution). The GM, WM and CSF probabilistic maps built and used in this study were calculated from 138 T_2_-weighted anatomical images, resulting in robust tissue class priors not only for VBM analysis but also for functional MRI and diffusion tensor imaging analysis in mice.

This comprehensive set of MRI compatible template and atlas for the mouse brain, allows a unified and standardized analysis of multimodal mouse brain MRI data and paves the way for the development of multicentric preclinical studies. Indeed, animal models deliver crucial information for the understanding of brain structure and function both in healthy and pathological conditions. Our template and mouse brain atlas were conceived to bridge the gap between basic and clinical neurosciences by providing to the preclinical neuroimaging community specific resources designed to be used in conjunction with the neuroinformatic tools and methodologies commonly used in human MRI studies. We anticipate that these resources will help neuroscientists to conduct their analyses of anatomical and functional datasets in a more standardized way, with the final goal of reaching more reproducible conclusions (https://www.nitrc.org/projects/tmbta_2019).

### Gestation and lactation periods induce strong but transient GMC hypertrophy

The establishment of accurate mouse MRI resources permit to study the variations of GMC longitudinally, *in vivo* and during the gestation and lactation periods in female mice. We observed that several brain regions became transiently hypertrophic during pregnancy or in the lactation period until weaning. A set of structures comprising the core of the maternal circuit – especially the mPOA and BNST – displayed long-lasting hypertrophy that started at late gestation, culminated during the first week of lactation, and then disappeared at weaning. The initial GMC increase observed during the gestation period probably reflects changes induced by hormonal priming^11^. Indeed, both the mPOA and BNST express a high number of steroid hormone and neuropeptide receptors^29,30^. These factors are well known to trigger significant plasticity changes within the core maternal circuitry that are necessary for the preparation and adaptation of the brain to motherhood^3,11^. From parturition, the GMC differences observed during the whole lactation period highlight that mPOA/BNST receives a variety of sensory inputs from the pups, integrates that information with the females’ endocrine status, and then projects to brain sites involved in socially-relevant motivation, affective state, and cognition^2,31^. Pup stimulation, electrolytic and neurotoxic lesions and local steroid hormone injections^32–35^ have been shown to modify the intrinsic activity of these nuclei and consequently responsible for motivation and expression of maternal behavior. The mPOA is engaged throughout the postpartum period but differentially according to the needs of the developing pups. It has been shown that neurons of the mPOA in late postpartum inhibits maternal responses allowing the changing expression and waning of maternal behavior across postpartum^10^. The sustained increase in GMC reported in late lactation could reflect the involvement of the mPOA to appropriately influence maternal behavior.

Our study highlights another set of brain structure that became hypertrophic only during the period from parturition to weaning – specifically, the PVN and arcuate nucleus of the hypothalamus. These changes illustrate the structural plasticity occurring in these regions. For instance, at parturition and during lactation oxytocin neurons of the PVN undergo dramatic neuronal, glial and synaptic changes such as an increase in size of the oxytocin neurons and an amplification of their synaptic input^36^. Oxytocin release at parturition facilitates the onset of maternal behavior by acting on the mPOA and is also important for maternal memory^37^. Finally, lesions of the PVN disrupt the onset of maternal behavior^38–40^. In the arcuate nucleus, dopaminergic cells are responsible for suckling induced prolactin release^41^ and neurons projecting to the arcuate nucleus are involved in the maintenance of maternal motivation^42^.

Additionally, we report changes in GMCs found in olfactory related structures (main olfactory bulb, piriform cortex), somatosensory areas and auditory areas which reflect the multisensory control of maternal behavior. This finding is in accordance with a functional MRI study performed in rats which demonstrates that pup suckling is associated with increased neuronal activity within the midbrain, striatum and cortical sensory areas (somatosensory, olfactory and auditory cortices)^9^.

GMC variations within the hippocampus and entorhinal cortex, highlight the role of two essential structures involved in learning and memory processing during the reproductive period. Our data support evidence that the hippocampus undergoes profound neural changes during lactation. Indeed, lactating females have elevated spine densities in the hippocampus^43^ and show significant dendritic remodeling in pyramidal neurons^44^. Changes in hippocampal neurogenesis occurs during lactation and may support the enhancement of spatial memory necessary to foraging behavior in lactating females^43,45–47^.

Finally, our study also revealed a hypertrophy of the agranular insular cortex, which has never been reported in this context. The agranular insular cortex is a laminar part of the insular cortex and can be considered as a hub structure linking large-scale brain systems^48^. Indeed, the insula receives direct thalamic and somatosensory afferents carrying sensitive stimulations. In addition to its sensory afferents, the insula displays structural connectivity with the limbic system (basolateral, lateral and central amygdalar nuclei) as well as with the BNST, mediodorsal nucleus of the thalamus, lateral hypothalamus and perirhinal and lateral entorhinal cortices^48^. The insula also connects brain regions implicated in motivation and reward, such as the nucleus accumbens and caudate putamen^48^. Hence, our findings and the current literature suggest that before and after birth, the insular cortex may integrate and combine information from both external and internal stimulation and act as a relay between higher cortical and subcortical structures. Together, our results describe the dynamics of neurophysiological adaptation occurring in the brain from the early gestation period to weaning, thereby ensuring efficient maternal behavior and, by extension, the development of the offspring.

### GMC modifications in the olfactory system at the end of the gestation predict the level of maternal behavior post-partum

All these transient modifications of GMC in parous animals indicate that various brain regions undergo considerable plasticity from birth to weaning. Then, we sought to determine whether inter-individual variations in maternal behavior were associated with similar variations of GMC. Based on their behavioral performance in the pup retrieval test, we used a k-means clustering strategy to divide maternal female mice into two groups displaying high *versus* low levels of maternal behavior. This analysis revealed that several transient and several long-lasting increases in GMCs were observed in the high maternal behavior group that were absent in the low maternal behavior group. Brain regions showing significant differences included the olfactory bulbs, somatosensory system, limbic system, especially the orbitofrontal area, and mnesic system, including the retrosplenial cortex, hippocampus and entorhinal area. Some of these structures are directly responsive to pup stimulation, and the observed dynamics may have been induced by mother-offspring interactions. For example, higher GMC values in the somatosensory cortex and olfactory bulbs in the high maternal behavior group potentially reflected increased suckling duration and proximity between the mother and pups, respectively.

Strikingly, differences in GMCs were detected in the entorhinal area, orbitofrontal area, olfactory bulb, hippocampus, retrosplenial area and primary somatosensory area before parturition. These findings suggest that the maturation of these structures, probably through hormone-dependent plasticity mechanisms, is a key determinant of the intensity of maternal behavior expressed during the lactation period. Using a ROC procedure, we found that GMC values of the entorhinal area and AOB at the end of gestation were significantly predictive of the maternal behavior postpartum. Interestingly, previous studies in mice reported an increase in cell proliferation during gestation in the subventricular zone, the neurogenic niche which provides newly generated neurons within the olfactory bulb (for review see ^13^). These adult-born olfactory neurons are fully responsive to pup odor exposure^49^ and are in part involved in some components of maternal behavior^50,51^.

Hence, the correlations observed between GMC values in AOB at the late gestation period and maternal behavior performances may suggest that impairments of neural plasticity of this olfactory region would induce maladaptive neuroendocrine processing of the maternal brain at the end of the gestation period impacting maternal behavior performance. Taken together, our data provide the first potential imaging-based predictive biomarkers of the quality of maternal behavior and suggest the key role of the maturation of the olfactory system at the end of the pregnancy in the development of adaptative maternal behavior in mice.

## Conclusion

Our study provides a new generation of neuroinformatic tools which will help basic neuroscientists to conduct structural and functional MRI investigations. Using these resources, we found that the development of the maternal brain is associated with substantial mesoscopic changes in critical regions. These modifications can be interpreted as cell size changes, neural or glial cell genesis/apoptosis, spine density or blood flow modifications^52–54^. As cellular and molecular plasticity events are key for the adaptation of the brain to motherhood, molecular, cellular and behavioral investigations must be performed to obtain a more precise view of the physiological mechanisms responsible for GMC variations.

## Methods

### Animals

Twenty-three female RjOrl:SWISS virgin mice (8 weeks old; 20-25 g; Janvier Laboratory, Le Genest-Saint-Isle, France) were maintained on a 12-h light/dark cycle with access to food (standard chow) and water *ad libitum*. Animals were acclimatized 6 per cage to the housing facility for 7 days prior to manipulation. Females were randomly divided into two groups: a parous group (n = 12), in which each female was exposed to a male (RjOrl:SWISS,8 weeks old; 20-25 g; Janvier Laboratory, Le Genest-Saint-Isle, France) for 5 days, became pregnant, and raised their offspring (litter size: 6 to 14 pups) until weaning at 21 days postpartum, and a control group (n = 11), in which virgin females were not exposed to male mice. Each parous female was individually housed after male exposure. Control females were housed together in a separate room from parous females.

The MRI protocol was optimized to keep mice anesthetized for 2 h during each of the six acquisitions. During lactation MRI acquisitions, pups were kept under a heat lamp. One week after birth, maternal behavior was assessed as described in the behavioral section. All experiments were conducted in accordance with the local research ethics committee (APAFIS #6626-201002281145814V1) and are reported in accordance with the ARRIVE guidelines.

### MRI acquisition

*In vivo* 3D MRI of the entire brain was performed three days before male exposure (baseline), at one week of gestation (early gestation), two days before the expected day of birth (late gestation), one week postpartum (early lactation), three weeks postpartum (late lactation) and two weeks after weaning (weaning). One female in the parous group and one female in the control group were scanned under similar conditions on the same day.

Mice were anesthetized using isoflurane (2.5%; induction in O_2_/air mixture 1:1) (TEM-SEGA, F-33600 Pessac, France) and then transferred and placed head first *procubitus* within an MRI-compatible cradle that incorporated a stereotaxic system dedicated for mouse head MRI, connected to a heater with circulating water to maintain body temperature and supplied with 1-2% isoflurane *via* a fitted mask. Respiration rate was recorded during all the experiments using an MRI-compatible monitoring system (PC-SAM model #1025; SA Instruments Inc., Stony Brook, NY, USA) and used to adjust the isoflurane rate to maintain a rate between 20 and 40 respirations per minute. After a recovery period of one hour, mouse returned to her pups. MRI studies were conducted at the Centre de Biophysique Moléculaire d’Orléans and were performed on a 7T/160 mm PharmaScan spectrometer (Bruker Biospin, Wissembourg, France) equipped with an actively shielded B-GA09 gradient set, with 90-mm inner diameter and 300-mT/m gradient intensity. A 23-mm inner diameter Bruker birdcage coil with a cradle dedicated to a mouse head was used. Data acquisitions were performed on an Advance III console running ParaVision 5.1 software. T_2_-weighted images were acquired using a 3D fast large-angle spin-echo (FLASE) sequence which allows 3D brain mapping with a high resolution in a suitable time for *in vivo* acquisition^55,56^. Thus, the sequence with echo time (TE) = 20◻ms, 1 repetition, acquisition matrix =160 × 140 × 95, and a field of view (FoV) of 19.2 × 16.8 × 11.4◻mm^3^, resulting in a final resolution of 120 μm isotropic voxels^55,56^. To obtain the FLASE sequence^55^, which is a specific sequence that is not included in the sequence package provided with ParaVision, the usual rapid acquisition with relaxation enhancement (RARE) spin-echo sequence was modified; in particular, the RARE-factor was fixed to 1 allowing a flip angle (FA) higher than 90° for the excitation pulse, while maintaining a 180° refocusing pulse^55^. Thus, T_2_-weighted images were obtained with a repetition time (TR) as short as 300◻ms, 10 times lower than that needed for a classical T_2_- weighted spin-echo sequence. The sequence was optimized for acquisition in 1 h 28 min, with an isotropic resolution of 120 μm, a TR of 300 ms, an TE of 20 ms, an excitation pulse (FA) of 115°, with 1 repetition and with a matrix of 160 x 140 x 95 corresponding to a FoV of 19.2 × 16.8 × 11.4 mm^3^ and contained the whole mouse brain.

### Maternal behavioral test

One week after birth, the maternal behavior of each female was evaluated using the pup retrieval test^57^. Briefly, three pups were removed from the nest and placed at three different corners within the home cage. The latency to retrieve each pup and the time spent licking the pups, crouching in the nest over the pups and performing nonmaternal behaviors such as self-grooming and digging were recorded over 15 min. Retrieval was defined as the animal picking up a pup and transporting it to the nest. Crouching was defined as the animal assuming the nursing posture. Nursing and licking were permitted whether they took place in the nest. All videos were analyzed using <u>BORIS software version 4.1.4</u>.

### K-means clustering

Clustering analysis of the behavioral data was performed using MATLAB Simulink 10b (The Mathworks, Inc., USA). Normality was verified and no outlier subjects were detected. To classify animals according to their maternal performance, a k-means clustering algorithm was used with crouching and digging times as behavioral markers. Digging was chosen because it is indicative of high maternal stress^28,58^. This algorithm iteratively grouped the animals by creating k initial centroids, assigning each animal to the closest centroid, iteratively re-calculating the centroids from the mean of its assigned animals and re-assigning the animals to each centroid until there were no more changes across iterations^59^. This clustering divided parous animals into a high maternal behavior group (with high crouching and low digging time) and low maternal behavior group (with low crouching and high digging time).

### Mouse brain template and atlas building

For the MRI protocol, we developed a brain template and an atlas from the <u>AMBMC brain template</u> and the <u>Allen Mouse Brain Common Coordinate Framework</u>, respectively (**Figure 1**). First, we down-sampled the AMBMC template and its associated atlases and the Allen Mouse Brain Atlas and its associated Nissl images to a suitable resolution for MRI analysis (60-μm isotropic resolution; **Figure 1**, step 1). Then, all images were manually aligned to the anterior commissure/posterior commissure (AC/PC) axis, and the center of the images was defined relative to the AC (**Figure 1**, step 2). The resulting template was then segmented into GM, WM and CSF probability maps using the unified segmentation approach^60^ of Statistical Parametric Mapping 8 (<u>SPM8</u>) and the mouse brain priors provided by the <u>SPMMouse toolbox</u> (**Figure 1**, step 3). In parallel, our T_2_- weighted anatomical images were realigned, coregistered, bias-corrected and normalized to our template. Using the SPMMouse toolbox, we also segmented the images as described above, and from these preprocessed images, we obtained a large set of 138 images for each tissue class (**Figure 1**, steps 4-7). From these images, we built population-specific GM, WM and CSF priors. To build these priors, for each tissue class, we applied a diffeomorphic anatomical registration using an exponentiated lie algebra (DARTEL) approach, which is an automated, unbiased, and nonlinear template-building algorithm^61^ (**Figure 1**, step 8). This new set of population-specific tissue priors was used for both atlas building and final VBM preprocessing.

To normalize the Allen Mouse Brain Atlas to our brain template, we used the associated Nissl-stained images because (1) Nissl staining corresponds to the GM prior in terms of histology, and (2) this image was already coregistered to the atlas (**Figure 1**, step 4). Therefore, we applied the segmentation function provided by SPM8 using the GM prior previously calculated from the Nissl image to generate the “Nissl2template” normalization matrix. We used this matrix to normalize the atlas to the template, while avoiding interpolation to maintain the label indices as integers (**Figure 1**, step 7). Then, a visual inspection of each normalized label was carried out to assess whether the normalization process modified the position and volume of the structure too much. When necessary, holes were filled and labels were redrawn according to Paxinos and Franklin’s atlas and using the <u>FreeSurfer</u> package. Finally, the olfactory bulbs and hind brain regions were completed, the corpus collosum and ventricles were drawn from the WM and CSF priors, and the cerebellum labels were replaced by the AMBMC cerebellum labels, which are more accurate. Finally, the atlas image was symmetrized (left-right). Our mouse brain template, priors and atlas were normalized within the same space and with the same final resolution (60-μm isotropic resolution), resulting in our final mouse brain atlas composed of a mosaic of 1320 ROIs covering the entire brain (**Supplemental Video 1**).

### VBM data preprocessing

Previously preprocessed normalized T_2_-weighted data were segmented into GM, WM, and CSF within SPM8 using the images of the population-specific priors (**Figure 1**, step 9). Then, to produce a more accurate registration within each mouse as well as across all mice, a longitudinal VBM analysis was applied using the strategy described by Asami *et al*^62^. First, a *subject-specific* template was created by the DARTEL algorithm using the previous tissue class images (*i.e.*, GM, WM, and CSF maps) obtained from each mouse for the six time points. The DARTEL procedure releases *individual-specific* flow field maps, permitting the application of diffeomorphic normalization on images of each tissue class to spatially normalize each time point on a *subject-specific* template space. Each normalized tissue class image was modulated by the Jacobian determinant to account for the expansion and/or contraction of brain regions over time. Then, a *population-specific* template was created by the DARTEL algorithm using all *subject-specific* templates of the tissue class images. Here, the DARTEL procedure releases *population-specific* flow field maps, permitting the application of diffeomorphic normalization of each animal onto the images of each tissue class. Finally, tissue class images were modulated by the Jacobian determinant, and the final modulated GM images were spatially smoothed with an isotropic Gaussian kernel with a 3-mm full-width at half-maximum and convolved with GMC images to create GMC maps (**Figure 1**, steps 10-12).

### VBM statistics and analysis

SPM8 was used to reveal the temporal and regional changes in the GMC maps. A second-level SPM analysis comprising a flexible factorial model, which is equivalent to a 2×2 mixed-model ANOVA with group as the between-subject factor and time point as the within-subject factor, was used to compare the control *versus* the parous groups and the low *versu*s high maternal behavior groups^63^. The factors included in the analysis were subjects, group (control *versus* parous, or low maternal behavior *versus* high maternal behavior), and time points (baseline, early gestation, late gestation, early lactation, late gestation, and weaning). A brain mask was used to constrain the analysis within the brain. For each cluster, the significance of the peak voxel was set as *p* < 0.01 (t_(126)_ = 2.356, control *versus* parous; t_(60)_ = 2.39, low *versus* high maternal behavior), and the minimum cluster extent was set at 25 voxels. The results are presented on axial brain slice series generated by the <u>Xjview</u> SPM plugin. Corresponding surfacing results were produced with <u>BrainNet viewer 1.6</u>^64^, allowing the generation of both brain meshes and brain plots to visualize data and create supplemental videos.

### Postprocessing statistical analysis

Cluster peaks revealed by the flexible factorial analyses were identified using our atlas and an *in situ* procedure developed with MATLAB Simulink 10b (The Mathworks, USA). For each comparison, clusters were binarized, and the obtained masks were used to extract GMC values of corresponding regions from the GMC map using the REX plugin. GMC data and behavioral measurements were then compiled and analyzed using GraphPad Prism 6.02 software. GMCs were compared between groups and for each time point using a two-way ANOVA with repeated measures followed by a two-stage setup method of Benjamini, Kriegger and Yekutieli as recommended by the software. Maternal and nonmaternal behaviors of the low maternal and high maternal groups were compared using a multicomparison t-test with a false discovery rate (FDR) approach (Q = 1%). Correlation analyses were performed using a parametric two-tailed Pearson test. Specificity and selectivity analyses were performed using the ROC curve method. Statistical significance was defined as p < 0.05 (*) for these analyses.

## Abbreviations

AC-PC: anterior commissure-posterior commissure
AMBMC: Australian Mouse Brain Mapping Consortium
AOB: cacessory olfactory bulb
BNST: bed nucleus of the *stria terminalis*
CNS: central nervous system
CSF: cerebrospinal fluid
DARTEL: diffeomorphic anatomical registration using exponentiated lie algebra
df: degree of freedom
FA: flip angle
FLASE: fast large-angle spin-echo
FoV: field of view
MRI: magnetic resonance imaging
MOB: main olfactory bulb
mPOA: medial preoptic area
GM: gray matter
GMC: gray matter concentration
PVN: paraventricular nucleus of the hypothalamus
RARE: rapid acquisition with relaxation enhancement
ROC: receiver operating characteristic
ROI: region of interest
TE: echo time
TR: repetition time
VBM: voxel-based morphometry
WM: white matter

## Acknowledgments

The authors acknowledge the regional council of Centre Val-de-Loire for funding this research through the IMACERVOREPRO grant (convention 201500104011, 2015-2018) awarded to Matthieu Keller.

## Author Contributions

**D.A.B.** contributed to the atlas and template building, data analysis, drafted and revised the manuscript. **A.E.** contributed to the imaging sequence troubleshooting, data analysis and contributed to the critical revisions. **F.S.** contributed to MRI sequence programming, imaging protocol setting, data acquisitions and contributed to the analysis and critical revisions. **H.A.**, **W.M.**, **E.C.**, **M.M.**, **S.M.** contributed to the study conception and design and contributed to the critical revisions of the manuscript. **F.L.** contributed to raise the funding, study conception and design and contributed to the critical revisions of the manuscript. **M.K.**, is the principal investigator of the study, raised the funding, coordinated the project, revised and validated the manuscript.

## Supplementary Data

**Table S1.**
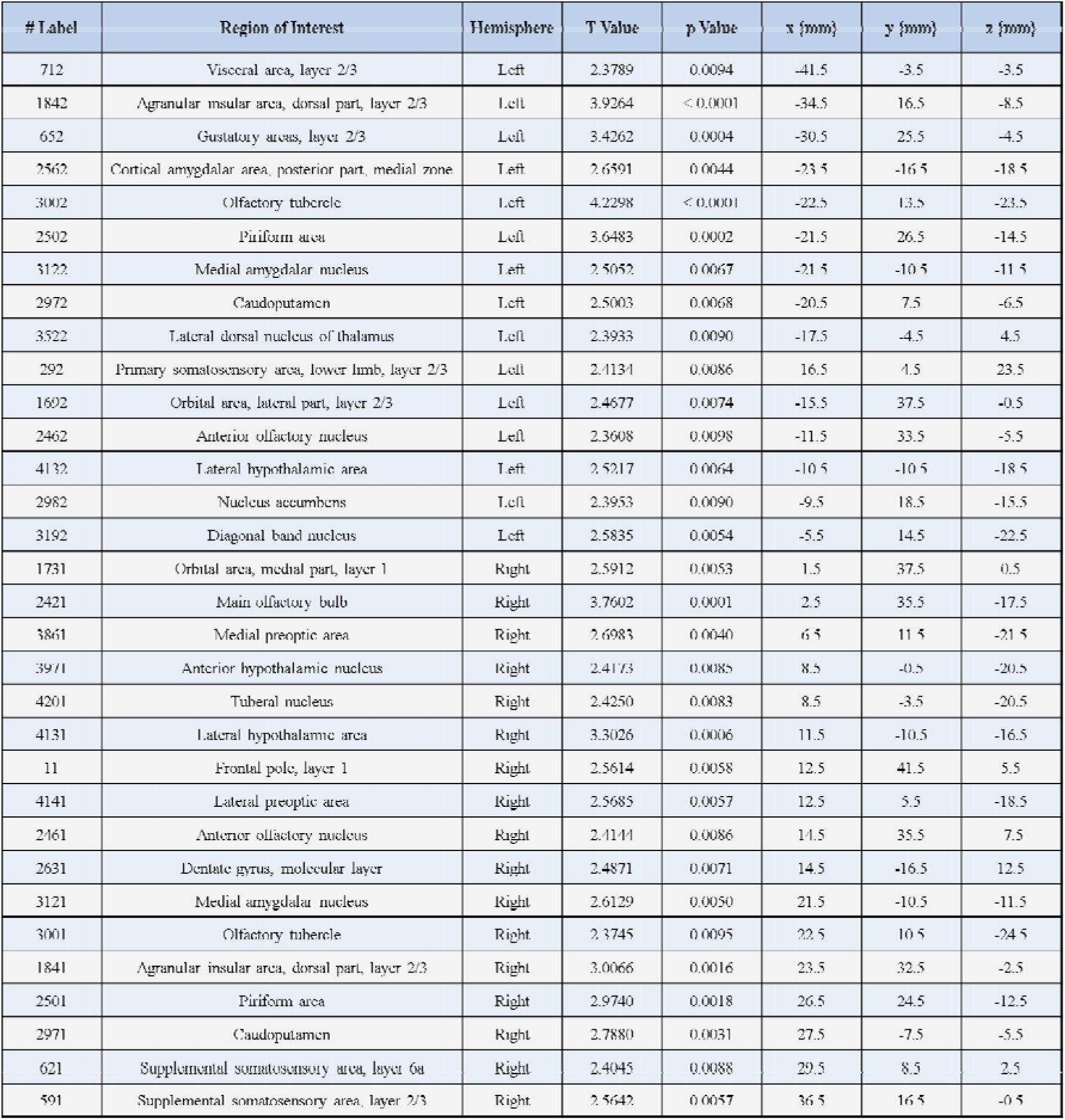
Local variation in gray matter concentration between control and parous mice at the end of the gestation period. SPM flexible factorial analysis revealed an interaction between the control and parous groups at the late gestation time point.

**Table S2.**
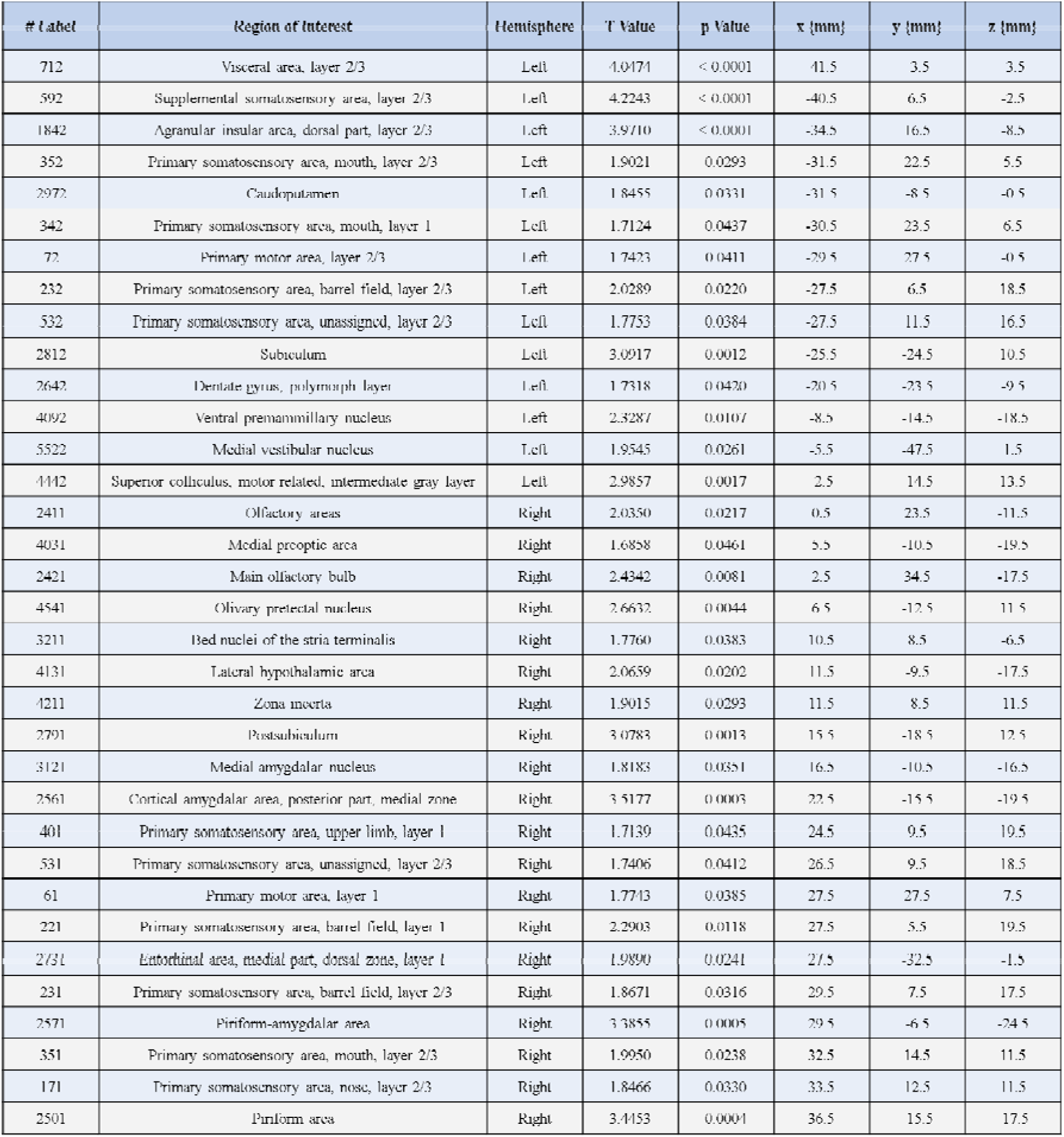
Local variation in gray matter concentration between control and parous mice at the beginning of the lactation period. SPM flexible factorial analysis revealed an interaction between the control and parous groups at the early lactation time point.

**Table S3.**
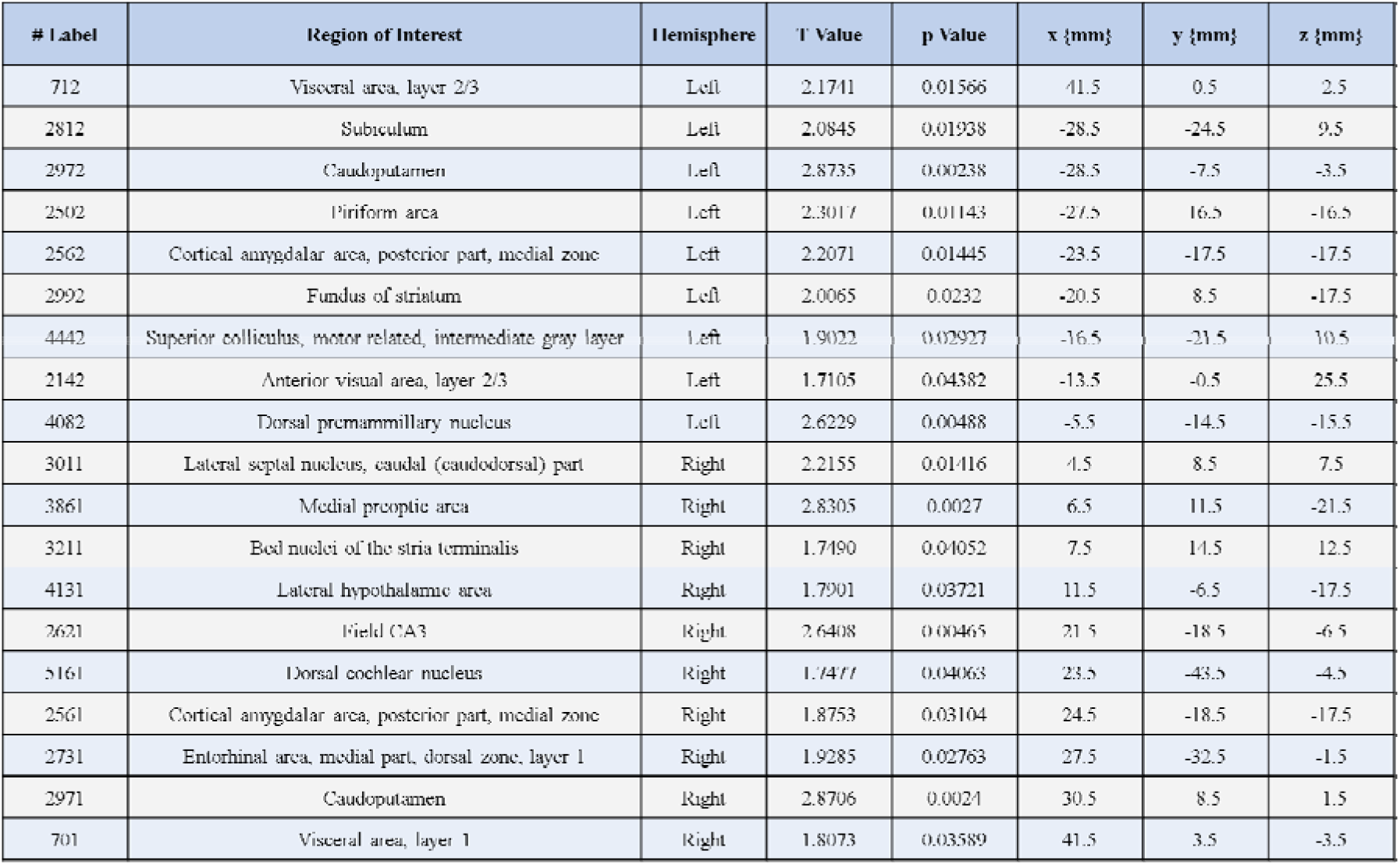
Local variation in gray matter concentration between control and parous mice at the end of the lactation period. SPM flexible factorial analysis revealed an interaction between the control and parous groups at the late lactation time point.

**Table S4.**
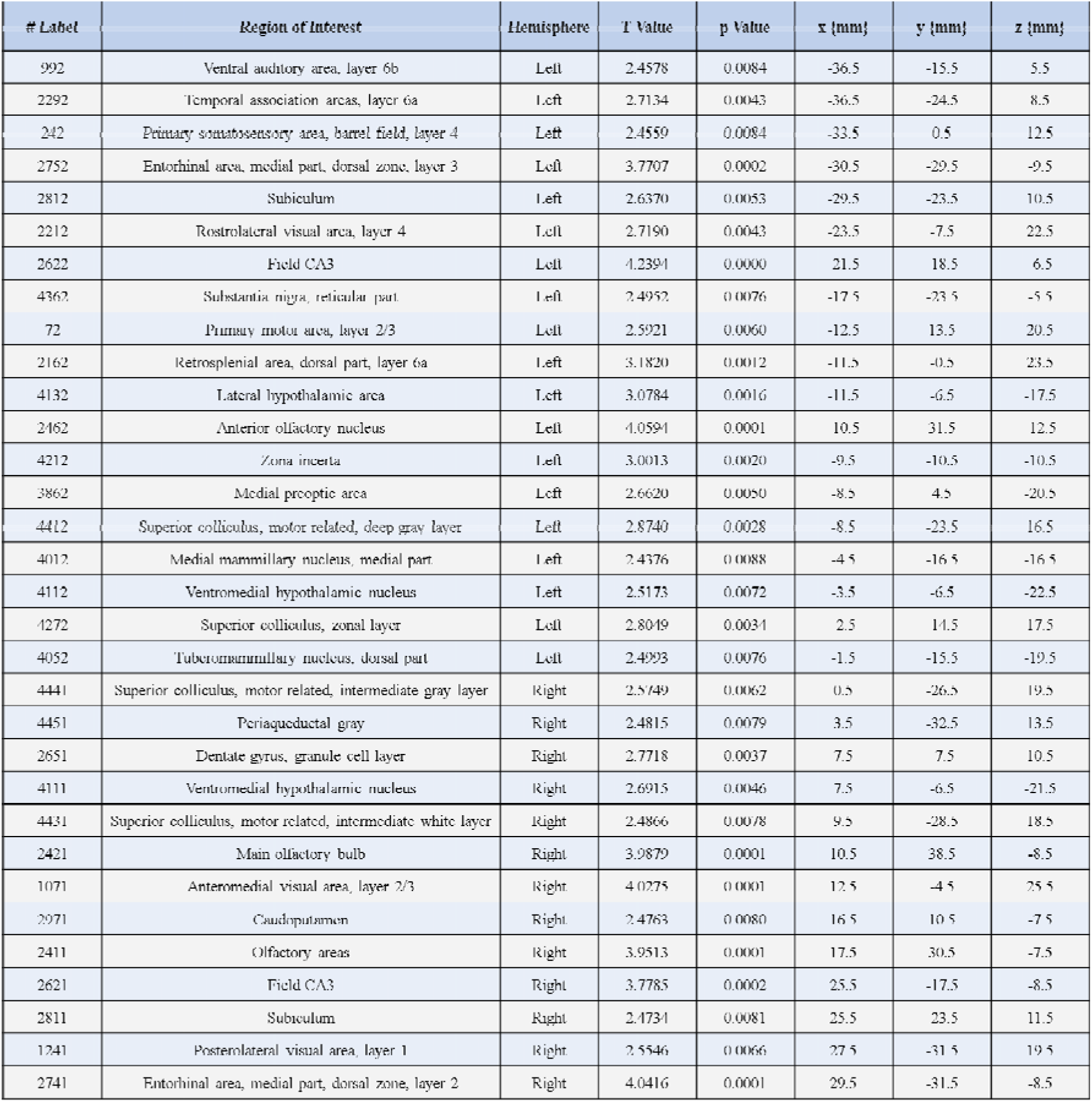
Local variation in gray matter concentration between low maternal behavior parous mice and high maternal behavior parous mice at the end of the gestation period. SPM flexible factorial analysis revealed an interaction between the low *versus* high maternal behavior groups at the late gestation time point.

**Table S5.** Local variation in gray matter concentration between low maternal behavior parous mice and high maternal behavior parous mice at the beginning of the lactation period. SPM flexible factorial analysis revealed an interaction between the low *versus* high maternal behavior groups at the early lactation time point.

| # Label | Region of Interest | Hemisphere | T Value | p Value | x {mm} | y {mm} | z {mm} |
| --- | --- | --- | --- | --- | --- | --- | --- |
| 2732 | Entorhinal area, medial part, dorsal zone, layer 1 | Left | 2.5882 | 0.0060 | 28.5 | 33.5 | 1.5 |
| 2632 | Dentate gyrus, molecular layer | Left | 2.5173 | 0.0072 | -20.5 | -16.5 | 6.5 |
| 412 | Primary somatosensory area, upper limb, layer 2/3 | Left | 3.1161 | 0.0014 | -19.5 | 14.5 | 18.5 |
| 2602 | Field CA1 | Left | 2.5276 | 0.0070 | -19.5 | -11.5 | 17.5 |
| 122 | Secondary motor area, layer 2/3 | Left | 2.8890 | 0.0027 | -15.5 | 24.5 | 15.5 |
| 2422 | Main olfactory bulb | Left | 3.5084 | 0.0004 | -8.5 | 45.5 | 6.5 |
| 3652 | Parafascicular nucleus | Left | 3.0525 | 0.0017 | -3.5 | -12.5 | -1.5 |
| 2672 | Induseum griseum | Left | 2.4662 | 0.0082 | -1.5 | 19.5 | 5.5 |
| 1482 | Anterior cingulate area, dorsal part, layer 1 | Left | 2.4675 | 0.0082 | -0.5 | 21.5 | 14.5 |
| 2082 | Retrosplenial area, ventral part, layer 1 | Left | 2.3913 | 0.0099 | -0.5 | -3.5 | 23.5 |
| 2421 | Main olfactory bulb | Right | 2.7201 | 0.0043 | 2.5 | 43.5 | -14.5 |
| 3651 | Parafascicular nucleus | Right | 2.9370 | 0.0023 | 6.5 | 12.5 | 0.5 |
| 2411 | Olfactory areas | Right | 3.2967 | 0.0008 | 8.5 | 42.5 | 4.5 |
| 2601 | Field CA1 | Right | 3.6131 | 0.0003 | 18.5 | -9.5 | 17.5 |
| 381 | Primary somatosensory area, mouth, layer 6a | Right | 3.1451 | 0.0013 | 24.5 | 14.5 | 5.5 |
| 2821 | Prosubiculum | Right | 2.9853 | 0.0020 | 24.5 | -19.5 | 14.5 |
| 2691 | Entorhinal area, lateral part, layer 2 | Right | 3.0830 | 0.0015 | 38.5 | 29.5 | 5.5 |

